# Male and female reproductive fitness costs of an immune response in natural populations

**DOI:** 10.1101/2020.07.10.197210

**Authors:** Stephen P. De Lisle, Daniel I. Bolnick

## Abstract

Parasites can mediate host fitness both directly, via effects on survival and reproduction, or indirectly by inducing host immune defense with costly side-effects. The evolution of immune defense is determined by a complex interplay of costs and benefits of parasite infection and immune response, all of which may differ for male and female hosts in sexual lineages. Here, we examine fitness costs associated with an inducible immune defense in a fish-cestode host-parasite system. Cestode infection induces peritoneal fibrosis in threespine stickleback (*Gasterosteus aculeatus*), constraining cestode growth and sometimes encasing and killing the parasite. Surveying two wild populations of stickleback, we confirm that the presence of fibrosis scar tissue is associated with reduced parasite burden in both male and female fish. However, fibrotic fish had lower foraging success and reproductive fitness (reduced female egg production and male nesting success), indicating strong costs of the lingering immunopathology. We show that these substantial sexually-concordant fitness effects of immune response act to align multivariate selection across the sexes, masking the signature of sexual antagonism that acted on morphology alone. Although both sexes experienced costs of fibrosis, the net impacts are unequal because in the two study populations females had higher cestode exposure. To evaluate whether this difference in risk should drive sex-specific immune strategies, we analyze a quantitative genetic model of host immune response to a trophically transmitted parasite. The model and empirical data illustrate how shared costs and benefits of immune response lead to shared evolutionary interests of male and female hosts, despite unequal infection risks across the sexes.

## Introduction

Organisms’ immune systems evolve to prevent or mitigate the fitness costs that parasites impose on their hosts. But, these immune systems can impose their own costs by consuming energy (Sheldon and Verhulst 1996) or attacking the host’s own tissues (Billi et al. 2019). Consequently, parasite infection reduces host fitness both directly, via effects on survival and reproduction, and indirectly by inducing costly immune responses (Khan et al. 2017, Armour et al. 2020). Host immunity is adaptive when the benefits of reducing direct effects of infection outweigh the indirect costs of the immune response itself. In this way, host immune response can be viewed and studied as a life history trait (Sheldon and Verhulst 1996, Schmid-Hempel 2003), whose expression may trade off with different fitness components and whose evolution depends upon these tradeoffs (Svensson et al. 1998, Lochmiller and Deerenberg 2000). Thus rather than simply being maximized, host immunity is expected to evolve to optimize these tradeoffs (Boots and Bowers 2004, Viney et al. 2005, Tschirren and Richner 2006, Houston et al. 2007, Graham 2013).

Evolution of immune defense becomes potentially more complicated when hosts reproduce sexually. Due to divergent gamete investment strategies and pervasive differences in the strength of sexual selection, males and females have sex-specific life history priorities (Rolff 2002, Stoehr and Kokko 2006). Shifts in the relative importance of longevity versus success in a single reproductive bout can alter the relative benefits of investing in immunological traits and mounting a response to a parasite or pathogen (Williams 1966, Zuk and Stoehr 2002, Gipson and Hall 2016, Hall and Mideo 2018). These strategic differences have been invoked to explain sex differences in the intensity and side-effects of male versus female immune response. There is some empirical evidence suggesting costs of immunity may be sex-specific: for example in humans there are numerous examples of sex specific immunopathology, including heightened prevalence of autoimmune disease in females (Fish 2008, Billi et al. 2019) and greater severity and frequency of parasitic diseases in males (Klein 2000, Roberts et al. 2001). These differences in optimal immune investment can be further exaggerated because males and females often differ ecologically (Shine 1989). Differences in diet, habitat use, or activity levels can impart distinct risks of infection (Bolnick et al. 2020), and consequently sex differences in parasite infection rates in wild animal populations are commonplace but variable (Poulin 1996, Schalk and Forbs 1997, McCurdy et al. 1998, Sheridan et al. 2000, Zuk 2009). These differences in life history and exposure suggest that sexes might benefit from evolving sex specific intensities of immune response, or different means of mitigating costs.

To test this proposition, we need studies that quantify the fitness benefits and costs of immune responses in nature, for both sexes. Are males and females equally at risk of parasite exposure? Do they initiate similarly strong immune responses? Are those immune responses equally beneficial by reducing parasite load? And, do both sexes experience equivalent costs of initiating these immune responses? Or, would differences at any of these stages lead to the evolution of sex-specific immune optima? Determining the causes of variation in sex-specific immunity and parasite infection is a major challenge in evolutionary ecology.

Here we explore the component fitness costs and benefits of immune response to parasite infection in threespine stickleback (*Gasterosteus aculeatus*), where we had reason to believe parasite exposure, infection costs, and immune costs might all differ between sexes. Of the suite of macroparasites that infect stickleback in postglacial lakes of the Northern hemisphere, the cestode tapeworm *Schistocephalus solidus* is one of the most abundant and causes high host morbidity in wild populations (Reimchen and Nosil 2001, Barber and Scharsack 2010, Stutz et al. 2014). Stickleback are an obligate intermediate host of *Schistocephalus solidus*, which are transmitted trophically when a stickleback consumes an infected cyclopoid copepod; sex differences in *Schistocephalus* infection rates appear to be related to differences in male and female diet (Reimchen and Nosil 2001). *Schistocephalus* complete their lifecycle when the host fish is consumed by a bird, which is their definitive host where the cestode mates. *S.solidus* is well known for its capacity to manipulate host color, behavior, and buoyancy to increase sticklebacks’ risk of avian predation. In addition to aiding host mortality, *Schistocephalus* infection is associated with reduced fecundity in females (Heins et al. 2010). Presumably to mitigate such fitness costs, some stickleback populations have evolved resistance to *Schistocephalus* infection, although other populations remain susceptible (Scharsack et al. 2007, Scharsack et al. 2016, Weber et al. 2017a, Weber et al. 2017b). In particular, fish in some populations develop peritoneal fibrosis in response to cestode infection, in which their body cavity and organs become engulfed in scar-like connective tissue (Weber et al in prep) similar to human fibrotic response during tissue repair (Mutsaers et al. 1997). In laboratory experimental infection trials, genotypes that initiated fibrosis were able to suppress cestode growth and occasionally trap and kill the parasite (Weber et al in prep). This fibrosis response is a deeply-conserved immune trait across ray-finned fishes, and can be induced by vaccination of a generic immune adjuvant (alum; Vrtlilek and Bolnick in preparation). However, some stickleback populations have recently co-opted this ancient pathway to respond to perceived *S.solidus* infections (Hund et al. 2020). Although both males and females respond to infection with similar levels of fibrosis, laboratory infection experiments reveal that fibrosis is associated with reduced female reproduction (Weber et al in prep). The web of fibrotic scar tissue might place an upper limit on ovary expansion. However, male reproduction does not rely on gonad enlargement and so we predicted that the costs of fibrosis may be sex-specific.

Two important questions in this stickleback-cestode system remain unanswered: why do populations vary in fibrosis immune response, and why do the sexes vary in cestode infection rates (Reimchen and Nosil 2001) but not in their innate immune response (Hund et al. 2020)? We predict that variation in fibrosis immune response across populations and cestode infection rates across sexes may be the result of differences in the cost/benefit trade-offs that dictate the optimal immune strategy. To test this prediction, we surveyed two wild stickleback populations that naturally differ in parasite infection rates. Both populations exhibit moderate rates of both infection and fibrosis. Crucially, there is an imperfect association between infection and fibrosis in both populations: not all individuals initiate fibrosis when infected. Also, fibrosis persists at least 3 months after the parasite is eliminated, so we find individuals with fibrosis but no surviving infection. These facts allow us to statistically separate the costs of infection from the costs of fibrosis, in both populations. We find massive fitness effects of both parasite infection and of the fibrotic immune response, in both sexes and across both populations. We show that these concordant fitness effects act to align otherwise-antagonistic multivariate selection across the sexes. These results leave us with a puzzle: there are persistent sex differences in infection rates, but no corresponding difference in the probability of fibrosis or its costs. To explain this apparent contradiction, we construct and analyze an optimality model of immune response evolution. Our analysis indicates that immune response optima may often be shared across sexes that differ in parasite encounter/infection rates when costs of infection and immune response are concordant and high.

## Methods

### Fish capture and measurement

We captured adult threespine stickleback from two lakes (Boot Lake and Roselle Lake), on Vancouver Island, British Columbia, Canada, between June 2-8 2019. These lakes were chosen because *Schistocephalus* infection and fibrosis response are observed in both, although to substantially different degrees; fish in Roselle exhibit a strong fibrotic immune response yet relatively low cestode infection rates, while fish in Boot exhibit much higher infection rates and lower rates of fibrosis (Stutz et al. 2014, Weber et al. 2017b, Hund et al. 2020). As noted above, the presence of both fibrosis and infection, imperfectly correlated, allows us to statistically partition their effects on measures of male and female reproductive success.

To compare reproductive success for males with versus without fibrosis (or, cestode infection), we compared the rates of fibrosis (infection) in randomly caught males, versus males that had successfully nested. We snorkeled in the littoral zone to search for nesting males; we identified nesting males based on their behavior (territory defense) and presence of a nest (e.g. arrangement of vegetative debris). Males exhibiting these qualities were observed until egg fanning behavior, or hatched fry, were observed, upon which the male was deemed to be a successful nester and was captured, immediately euthanized, and placed on ice and frozen. During this period we also placed minnow traps nearby to capture a random sample of males and females; this allowed us to obtain a random sample of unmated males for comparison to those that were successfully defending nests. Thus we obtained data required to measure total sexual selection acting via variance in male mating success (defining sexual selection sunsu Arnold and Wade 1984b, Arnold and Wade 1984a). Traps were placed at varying depths and distance from shore, and we avoided placing traps directly in areas of high nest density so most trapped males are unlikely to be nesting. Trapped fish were euthanized and frozen. We sampled both lakes until at least 50 nesting males had been captured. This number was chosen to avoid excessive impact on the populations studied while still presenting a reasonable sample size; we note however that this sampling design precludes accurate estimation of population mean mating rate because we have controlled the number of nesting males captured. We also retained a random sample of trapped females in each lake, using ovary mass as a metric of reproductive stage.

All fish were kept frozen until later laboratory dissection, upon which fish were thawed, measured, and dissected. We measured seven external morphological traits: standard length, head length (measured from the snout to the distal end of the operculum), snout length (measured from the snout to the orbital), eye width, body depth, body width at the pelvic girdle, and middle spine length. These traits were measured to two decimal places with digital calipers. We then dissected the left gills and counted gill raker number, in addition to photographing the gill rakers under a dissection microscope at fixed magnification to measure length of the longest raker. Following these measurements fish were dissected and all *Schistocephalus* counted and weighed. We note that *Schistocephalus* was the only abundant macroparasite found in our sample; we also examined internal organs, eyes, and the digestive tract for other parasite taxa. Gonads were removed and weighed. Stomach contents were removed and identified to (at least) order for all individuals, with the exception of samples from nesting males from Roselle, whose stomachs were lost during shipping.

We scored fibrosis on an ordinal scale, where a value of 0 corresponds to no apparent fibrosis (organs move freely), a value of 1 corresponds to fibrosis between organs, a value of 2 corresponds to fibrotic connection between organs and the peritoneal tissue, and a value of 3 corresponds to excessive fibrosis across the entire body cavity. This scale is modified from that developed to approximate the range of fibrotic variation seen in both laboratory studies and natural populations of stickleback (A. Hund & L. Fuess, personal communication), and has previously been shown to be highly repeatable between independent observers blind to experimental vaccination treatment (Goldzmid and Trinchieri 2012). A video example of these fibrosis levels is available (https://www.youtube.com/watch?v=yKvcRVCSpWI&feature=youtu.be).

### Statistical Analysis- Benefits of fibrosis

To evaluate the benefits of fibrosis, we used a generalized linear model to test whether *Schistocephalus* mass depends on host fibrosis score (treated as a continuous fixed effect which accommodates the ordinal nature of the variable), assuming exponential error. We avoided including fish body mass in this model because *Schistocephalus* mass is a direct component of total fish body mass and fibrosis is unrelated to mass, although results were unchanged when correcting for mass. Because our sample size for this analysis was limited to the subset of fish that were infected, we pooled lakes and sexes and did not model higher order interactions. We used a separate linear model with infection probability as a binomial response (cestode present or absent) to host fibrosis score, lake, and host sex; interaction terms were not significant and were dropped.

### Statistical Analysis- Costs of fibrosis and infection

We considered the fitness costs of fibrosis and infection on male and female reproductive success (nesting or not; ovary mass). We assessed component fitness costs for females in Roselle lake using a linear model with ovary mass as a response, fibrosis score and infection status as fixed effects, and exponential error. Small sample size of gravid females precluded such an analysis for Boot fish. We obtained qualitatively equivalent conclusions including body mass in the model, although present the results without size correction. A caveat with our analysis of female ovary mass is that we cannot control for females that may have already laid a clutch and are in the early stages of developing a second. We repeated this analysis for males from Boot and Roselle lakes, using binomial nesting success as the response variable and fitting separate models for each lake. Male testes mass is less clearly related to reproductive success than is female ovary mass and so we focus on male nesting success.

The above analysis of male nesting success indicated strong sexual selection against fibrosis, and we were interested in if and how this selective force acts in conjunction with sexual selection on morphology. To do this, we estimated the bivariate fitness surface for morphology and fibrosis for males separately for Boot and Roselle lakes using binomial linear models with nesting success as the response variable and fibrosis score and score on the multivariate morphological selection gradient as fixed effects. Score on the morphological selection gradient was obtained as discriminant function scores calculated in discriminant function analysis of nesting success and morphology, performed separately for each lake; this is equivalent to calculating individual scores on the vector of multivariate directional sexual selection on morphology (Mitteroecker and Bookstein 2011). This approach essentially estimates individual score on the morphological sexual selection gradient and then uses this score as a predictor in a linear model with fibrosis as an additional predictor of fitness. This approach allowed us to 1) estimate the effects of fibrosis on fitness accounting for effects of morphological traits, where morphology has reduced dimensionality yielding increased power and 2) allowed us to plot the corresponding bivariate fitness surface. We obtained qualitatively equivalent conclusions on the importance of fibrosis using a full multivariate model; we explore such a multivariate model below (*Statistical Analysis - Alignment of multivariate selection across the sexes*).

To assess differences in diet, including potential costs of immune response and infection on resource acquisition, we used a series of uni- and multivariate linear models to test for associations between diet and fibrosis or infection. Because of the sparse multivariate nature of diet data (i.e., many zeros for rare taxa), we constructed our models to test three specific hypotheses, in all cases avoiding higher order interactions wherever appropriate. First, to evaluate variation in overall prey intake, we used a univariate linear model with total prey counts as the response and fibrosis, lake, and infection status as fixed effects. Prior studies have reported inconsistent sex-biased diet in stickleback (males often being more limnetic in most but not all lakes; Bolnick and Ballare 2020). This is relevant because cestodes are acquired by eating limnetic copepods. To assess differences across the sexes in diet composition we used a multivariate linear model where the response vector consisted of counts of each prey taxon in an individual’s diet. This model included sex, lake, and their interactions with prey taxon as fixed effects; in this model significant interactions with taxon type indicate differences in the taxonomic composition of diet across sexes or lakes. Finally, we used a similar multivariate model to test for diet differences between nesting and randomly sampled males in Boot lake, although as a caveat we cannot control for variation in time spent in traps before euthanasia that may affect measured diet. Multivariate models with infection status as a predictor failed to converge, likely due to the relatively small number of infected fish. For all models of diet content, in which the data are counts of individual prey items in a stomach, we treated the response variable(s) as Poisson distributed. For multivariate models with a response vector of counts of each prey taxa, we modeled covariance in counts across individual fish as the Cholesky parameterization of an unstructured covariance matrix.

### Statistical Analysis - Alignment of multivariate selection across the sexes

Our analysis of fitness costs of fibrosis suggested shared reproductive costs of fibrosis across the sexes, and we were interested in quantifying effects of fibrosis on the geometry of multivariate selection. To assess the effects of fibrosis immune response on alignment between male and female selection, we compared the orientation of multivariate selection in males and females estimated in Roselle lake. Small sample size of females in Boot lake precluded such an analysis for those fish. For this analysis, we re-fit our model of male morphological sexual selection using a glm with nesting success as a response and morphological traits as a predictor, fit an equivalent model for females with ovary mass as a response, and compared the orientation of male and female selection (assuming gaussian error for both for comparison). We fit models including all morphological traits, and also models with reduced dimensionality where we used the first three principal components of the correlation matrix of morphological traits (both sexes pooled). We repeated this with and without including fibrosis score in as a predictor in our model in order to assess the effects of fibrosis on the alignment of selection across the sexes. Note that the non-binary fitness component for females precluded discriminant analysis as performed before for males. We compared orientation by calculating the vector correlation between the normalized selection gradients, ***β*_m_*β*_f_**^T^, which provides an estimate of sexual antagonism on a scale of −1 to 1, where a value less than zero indicates selection acting on opposing directions across the sexes, while a value greater than zero indicates concordant selection. Note that this approach compares orientation only, and not magnitude of selection, and so avoids likely issues arising from use of different fitness components in estimation of ***β*_m_** and ***β*_f_**. We estimated the sampling distribution of this vector correlation by resampling (100,000x) each vector from a multivariate normal distribution centered at the original REML estimates of ***β*** with covariance equal to the covariance matrix of the fixed effects from the fitted linear model, calculating the vector correlation for each sample. This approach is a modification, focusing on fixed effects, of a similar resampling approach (Houle and Meyer 2015) used to estimate sampling distributions of arbitrary functions of parameter estimates of random effects.

We focus our selection analyses on absolute component fitness, rather than relative fitness, because our fitness component estimates precluded estimation of population mean component fitness. This is because, by targeting a minimum number of nesting males, we lack a meaningful estimate of population-wide mean nesting rate. In such a situation, use of an arbitrary estimate of mean fitness to relativize fitness can be misleading when comparing selection among groups (De Lisle and Svensson 2017). Moreover, a focus on absolute component fitness is also appropriate because our interest is in performance costs associated with immune traits, rather than potential evolutionary response per se (De Lisle and Svensson 2017).

All statistical analyses were performed in SAS/IML version 9.3 (Cary Institute, NC). Generalized linear models were fit by REML using the glimmix procedure. Raw data and complete script to reproduce all analyses and figures is provided in the supplemental material.

## Results

### Benefits of fibrosis

We sampled a total of 411 fish (Roselle, 156 M, 71 F; Boot, 154 M, 30F), of which 102 were nesting males (50 Roselle, 52 Boot) and 81 (16 Roselle, 65 Boot) were infected with *Schistocephalus* (Table S1). Probability of cestode infection varied across sexes and lakes, with higher (odds ratio 4.79) infection rates in females across both Boot and Roselle lakes, and higher (odds ratio 10.5) overall infection rates in Boot than Roselle lake (Sex effect, *F*_1,407_ = 21.06, *P* < 0.0001; Lake effect, *F*_1,407_ = 46.14, *P* < 0.0001; Figure 1). Across lakes and sexes, probability of cestode infection was reduced in fish with high levels of fibrosis (odds ratio of unit offset at the mean 0.649, *F*_1,407_ = 6.15, *P* = 0.0135; Figure 1), consistent with prior observations that fibrosis contributes to elimination of the infection then lingers afterwards (Weber et al. in prep, Hund et al. 2020). Of those fish infected, host fibrosis was associated with reduced cestode mass (*F*_1,76_ = 9.06, *P* = 0.0035; Figure 2), consistent with laboratory infection experiments (Weber et al. in prep). Thus, across sexes and lakes, fibrosis confers benefits in the form of both a reduction in cestode growth and a decrease in the probability of infection.

**Figure 1.**
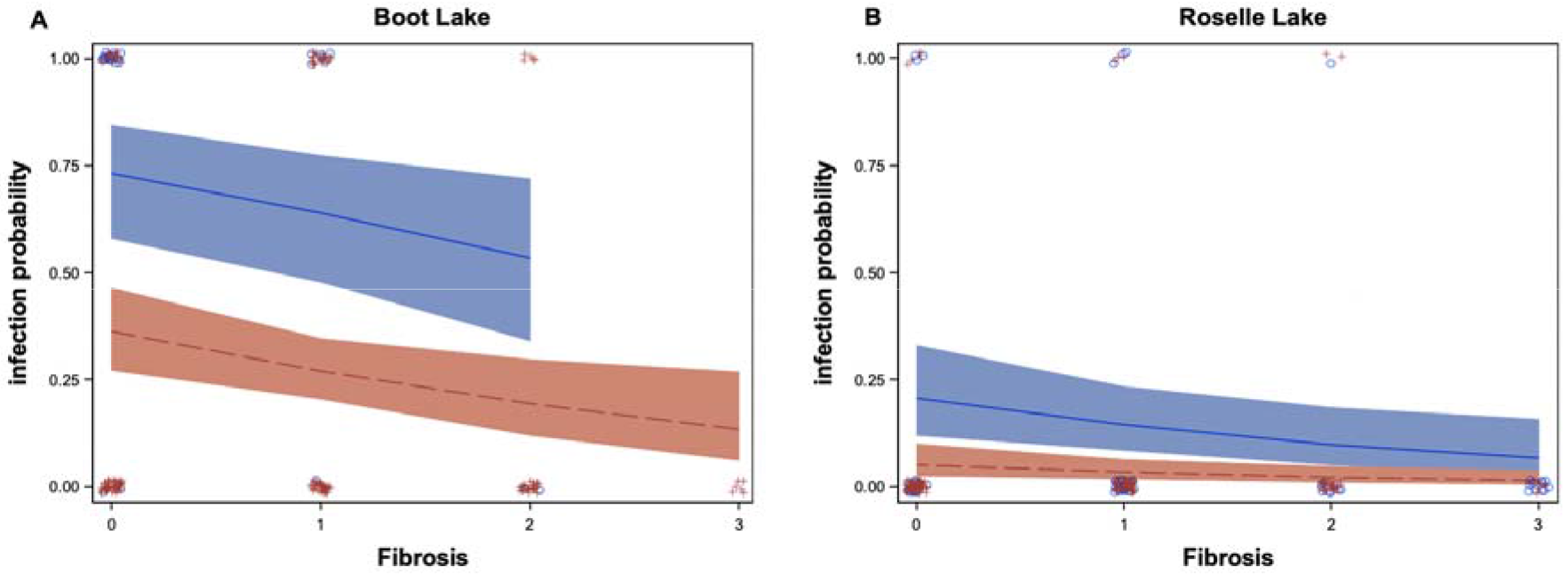
Fibrosis phenotype is associated with reduced probability of cestode infection across lakes and sexes. Model fit is from a generalized linear model with binomial error and lake, sex, and fibrosis as fixed effects. Interaction terms were not significant and were dropped; all main effects were significant (see text). Blue = females, red = males.

**Figure 2.**
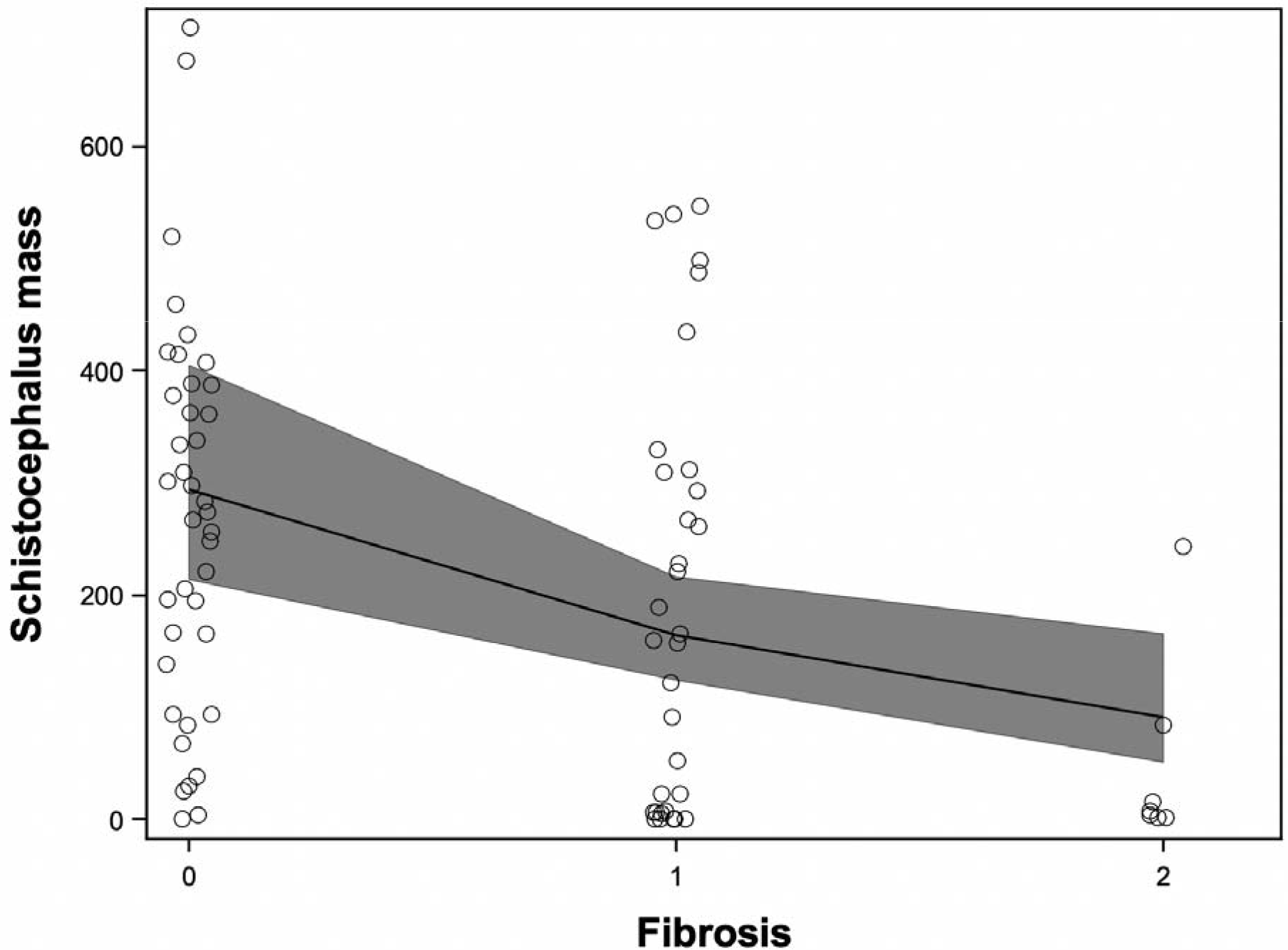
Fibrosis phenotype is associated with reduced cestode mass in infected individual hosts. *Schistocephalus* mass is the total mass of all worms found in the individual host. Model fit is from a generalized linear model with exponential error. Data are pooled across lakes and sexes; small sample sizes of infected individuals in lake x sex combinations make analysis of higher order interactions tenuous.

### Costs of fibrosis and infection

Countering these benefits, we also see evidence for costs of fibrosis. In both Boot and Roselle lakes, males defending active nests exhibited reduced fibrosis levels compared to randomly sampled males (Boot, odds ratio of unit offset at the mean 0.56, *F*_1,151_ = 6.87, *P* = 0.0097; Roselle, odds ratio of unit offset at the mean 0.41, *F*_1,153_ = 12.57, *P* = 0.0005) and reduced cestode infection in Boot lake (odds ratio 2.45, *F*_1,151_ = 4.3, *P* = 0.039; Roselle lake odds ratio 1.89, *F*_1,153_ = .57, *P* = 0.45). This is reflected in significant (Roselle, *F*_1,152_ = 6.24, *P* = 0.0135) or nearly significant (Boot, *F*_1,147_ = 3.52, *P* = 0.062) negative effects of fibrosis on mating probability in models including morphological discriminant function score, that is the vector of sexual selection on morphology, as a predictor of mating success (*P* < 0.0001 for both lakes). Thus, for both lakes, nesting success was determined by the independent effects of ecomorphology and fibrosis (Figure 3).

**Figure 3.**
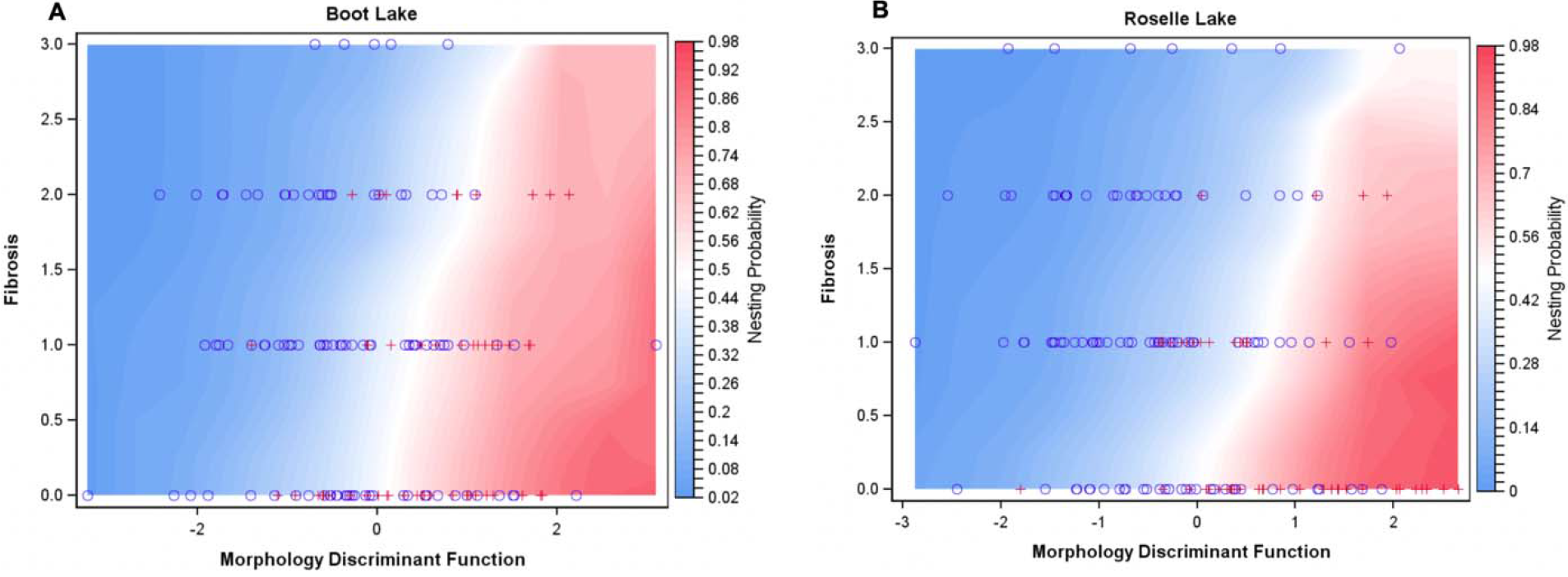
Sexual selection on fibrosis and morphology. Heat maps show male nesting probability as a function of fibrosis and sexually-selected multivariate morphology. The x axis is the discriminant function vector of morphometric traits that defines the direction of multivariate directional sexual selection on morphology alone, estimated separately for each lake. In both lakes, sexual selection acts against fibrosis independently of selection on size and shape. See text for details.

Although we lack data on mating success for females, we did obtain data on ovary mass, which is expected to be directly related to fecundity. For females in Roselle lake, we found that fibrosis was associated with a significant reduction in ovary mass (*F*_1,67_ = 10.65, *P* = 0.0017; Figure 4) while controlling for infection; ovary mass was also significantly reduced in cestode-infected females (*F*_1,67_ = 4.92, *P* = 0.03). Notably, the fibrosis response explained more variance in female ovary mass than the cestode infection itself explained. These results remained qualitatively equivalent when including body length as a covariate in the model. From Boot Lake we trapped too few gravid females to warrant analysis. Effect sizes for costs and benefits of fibrosis in both sexes are also presented below (see ‘A model of immune response evolution’).

**Figure 4.**
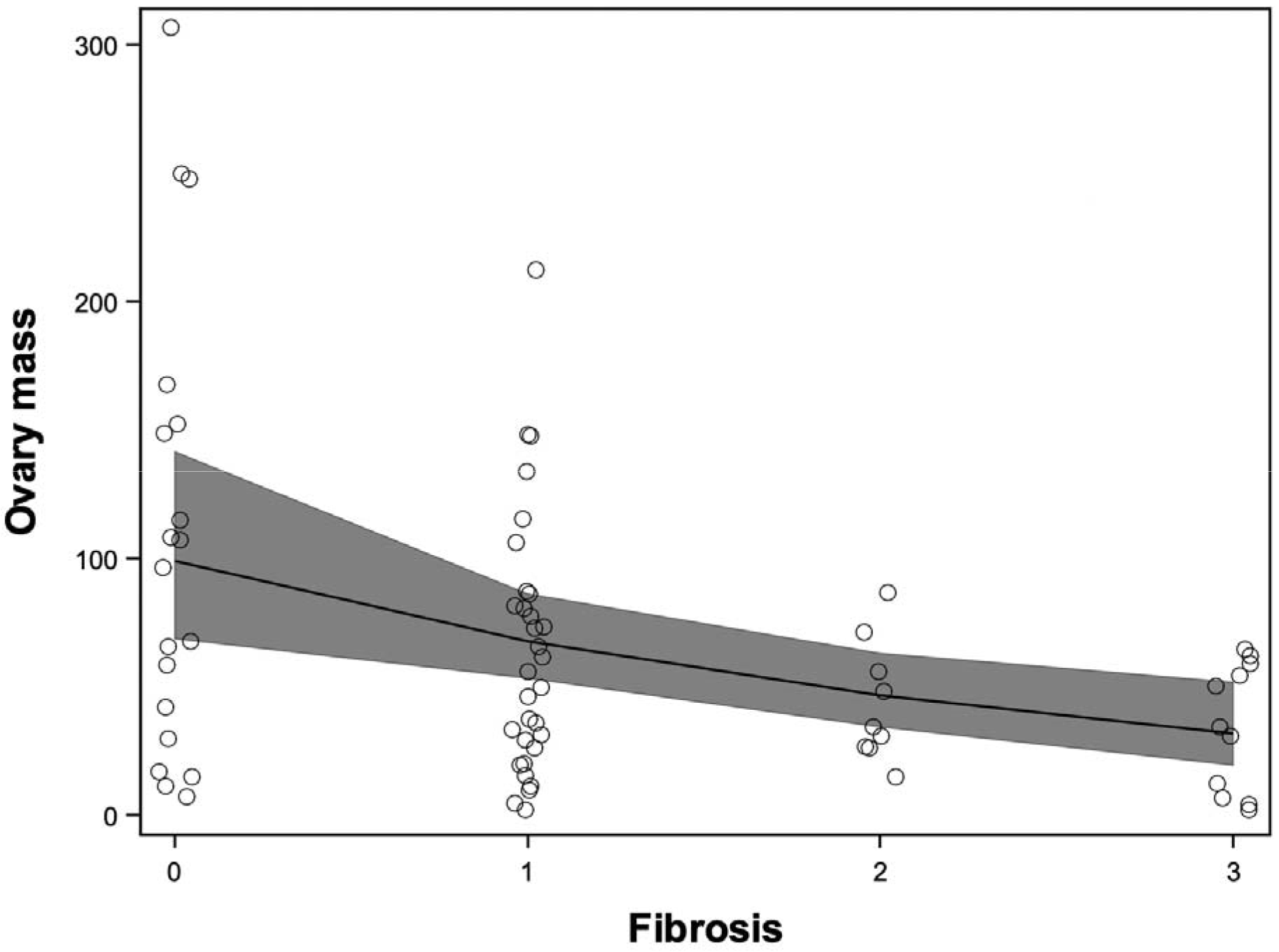
Fibrosis is associated with reduced female ovary mass in Roselle lake. Model fit is from a generalized linear model with exponential error. The small sample size of gravid females in Boot precluded analysis of female fitness effects of fibrosis.

In our analysis of diet content, we found individuals with high fibrosis had fewer total prey items in their stomach (*F*_1,341_ = 7.75, *P* = 0.0057; Figure 5A) controlling for lake effects (*F*_1,341_ = 43.11, *P* < 0.0001); this model indicated no difference in total prey items in infected versus uninfected individuals (*F*_1,341_ = .79, *P* = 0.37). In a multivariate model, we find no evidence of sex differences in the total number of prey items in an individual’s diet (sex effect, *F*_1,343_ = 0.06, *P* = 0.81), although the taxonomic content of individual diet varied with sex (sex*prey taxon effect, *F*_7,2393_ = 2.39, *P* = 0.019) while controlling for lake effects (lake*prey taxon effect, *F*_7,2393_ = 4.04, *P* = 0.0002). This sex effect was primarily driven by differences in the abundance of dipteran larvae, fish eggs, and zooplankton (Figure 5B), with females tending to have more limnetic prey. For the subsample of males from Boot lake, for which we had diet data for both nesting and non-nesting males, we found a main effect of nesting status in a multivariate model (*F*_1,145_ = 7.15, *P* = 0.0084) with nesting males having more prey items in their stomachs than a random sample of males in the population (Figure 5C). We found no evidence of a difference in the relative amounts of prey taxa in the diet of nesting and non-nesting males (mate status*prey taxa effect, *F*_3,431_ = 7.15, *P* = 0.35).

**Figure 5.**
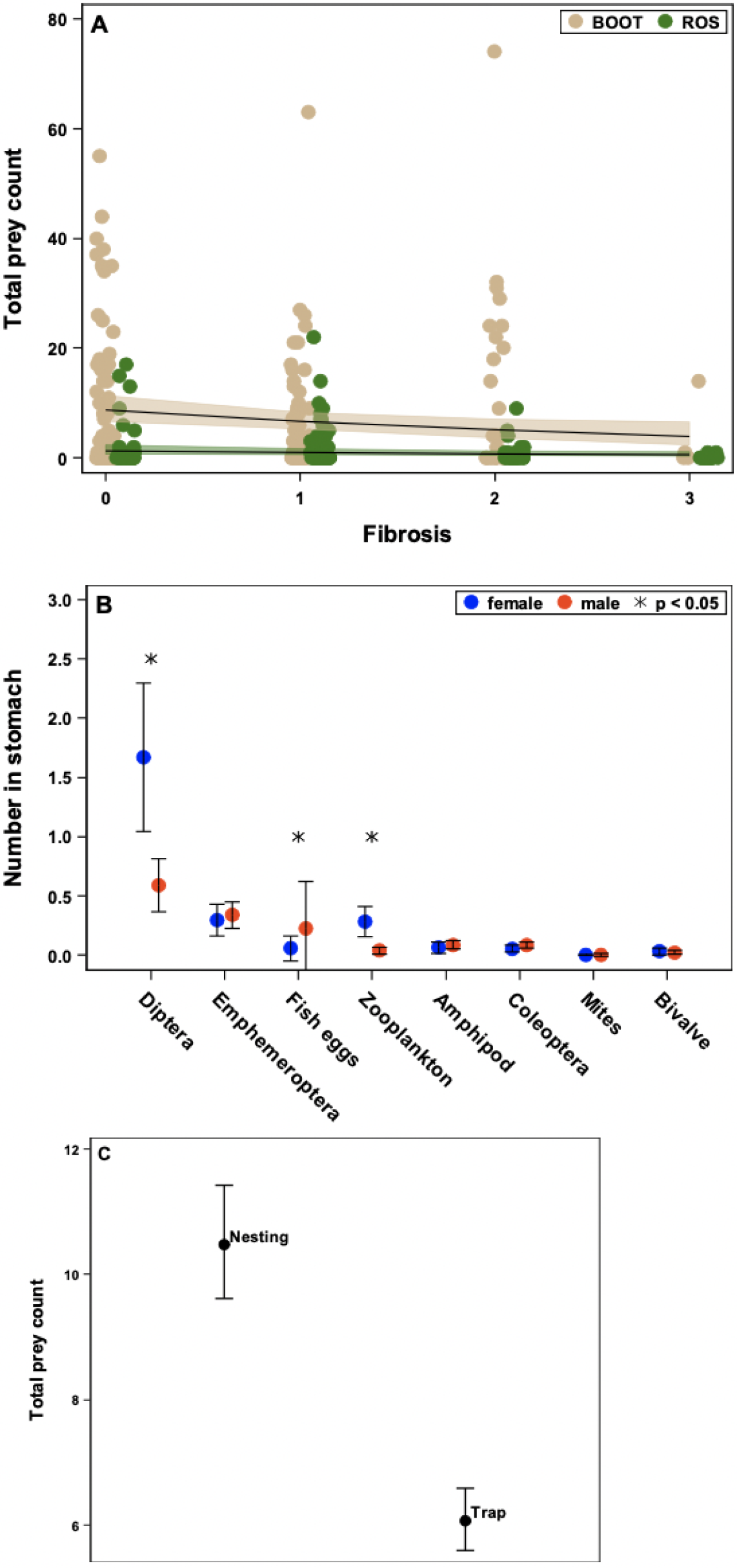
Individual diet varies with fibrosis, sex, and mating status. Across both lakes, fibrosis was associated with a reduction in the total number of prey items found in an individual’s stomach (panel A). Controlling for differences among lakes, males and females differed in their multivariate diet content, and this effect was driven by females consuming more dipteran larvae and zooplankton and males consuming more fish eggs (panel B; least square means and standard errors). In males from Boot lake, from which stomach contents of nesting males were available, we find that successfully nesting males had consumed more total prey items than a random sample of males captured in minnow traps (panel C; least square means and 95% CIs). From univariate (A and C) and multivariate (B) linear models with Poisson error.

### Alignment of multivariate selection

Given that fibrosis appears to confer similar component-fitness costs in males and females, we can examine the effects of fibrosis on geometric alignment of multivariate selection across the sexes. We find weak or even antagonistic alignment between multivariate selection on morphological traits in males and females (Figure 6), with evidence of sexually antagonistic selection on body shape independent of body size. This is reflected in a negative estimate of the correlation between male and female multivariate selection gradients (Figure 6B). However, including fibrosis as a predictor of component fitness results in a geometric alignment between male and female multivariate selection (Figure 6, A-C). Note that we avoid formal statistical comparison of male and female selection gradients because they were estimated in separate models using different fitness components.

**Figure 6.**
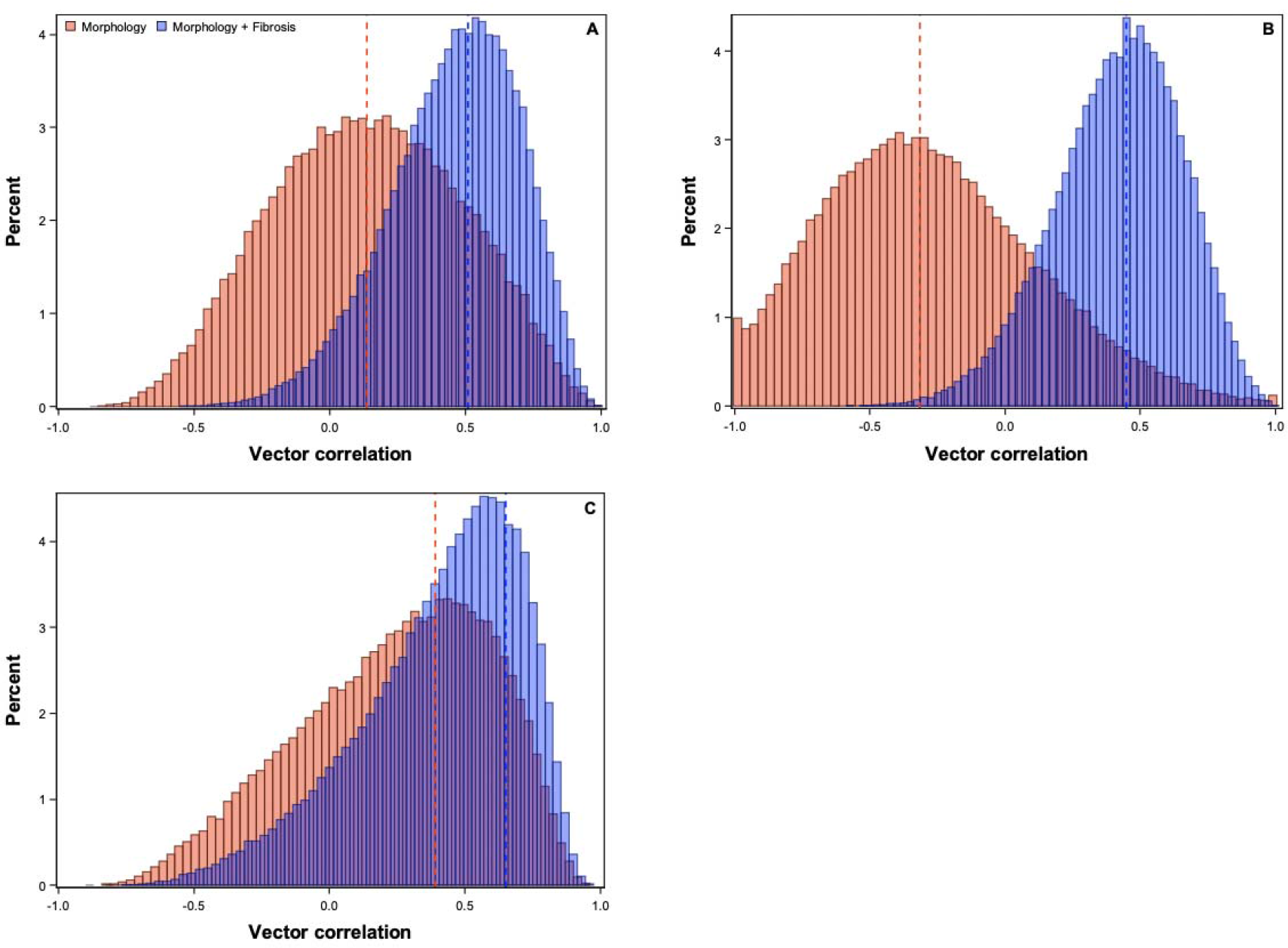
Sexually-concordant effects of fibrosis on reproductive fitness act to align multivariate selection across the sexes in Roselle lake. Histograms represent the empirically constructed sampling distributions of the vector correlation between male and female multivariate selection gradients estimated in separate linear models, when gradients are estimated using morphological traits alone (red) or including fibrosis (blue). Panel A shows this contrast in an analysis using the first three morphological principle components (i.e., size and two dimensions of shape that in total capture 85% of the phenotypic variation). Panel B shows the contrast in an analysis using PC2 and PC3, showing sexually-antagonistic selection on shape traits alone. Panel C shows the contrast when all raw morphological traits are modelled. Dashed lines represent the correlation estimated from the original REML estimates. Histograms are correlations calculated from gradient vectors obtained from resampling (100,000x) from a multivariate normal distribution centered on the original estimates and covariance from the estimated covariance matrices of the fixed effects.

## A model of immune response evolution

Our empirical results indicate substantial sexually-concordant fitness effects of fibrosis. We also find sex differences in cestode infection rates in the two populations we surveyed, and sex-specific infection rates are commonplace but variable across populations in Western Canada (Reimchen and Nosil 2001; Bolnick unpublished data). Yet, males and females do not appear to differ in fibrosis response (Hund et al. 2020, Weber et al in prep). These results are somewhat paradoxical because intuition suggests substantial sex differences in infection rates may be expected to lead to the evolution of sex-specific immune response. Here we construct and analyze a simple model of immune response evolution, in order to reconcile these results and illuminate how shared fitness costs of immunity may in part determine the optimum immune response in males and females.

We consider how selection acts on an underlying latent trait, the sensitivity of immune response to parasite exposure. Proximately, such a latent trait reflects mobilization of the immune system in response to a parasite infection/exposure. We seek to identify the rules that would determine the optimum sensitivity of immune response to parasite exposure and infection.

We define individual fitness as a function of expressed immune response and infection status,

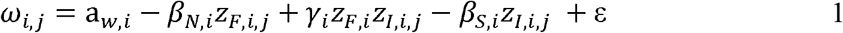

where *ω_i,j_* is the fitness of individual *j* of sex *i*. In this linear equation a*_w_* is an intercept (keeping in mind mean male and female fitness will be equal in sexual populations with a Fisherian sex ratio; Kokko and Jennions 2003), *β_N_* is the natural selection cost of an immune response trait, fibrosis *z_F_*, and *β_S_* is the cost associated with being infected, a trait termed *z_I_*. Definitions of all parameters in our model are presented in Table 1. The coefficient *γ_i_* represents the fitness benefit of an immune response, which manifests depending on both infection status and immune response phenotypes. Alternatively, we could define an alternative but equivalent algebraic expression for *γ_i_* where it is defined as a multiplicative function of *β_S_*; we retain the formulation in equation 1 because it is more readily estimable from data. The traits *z_F_* and *z_I_* can be quantitative on a latent scale (i.e., describing probability of immune expression or parasite infection), continuous, or binary.

**Table 1.**
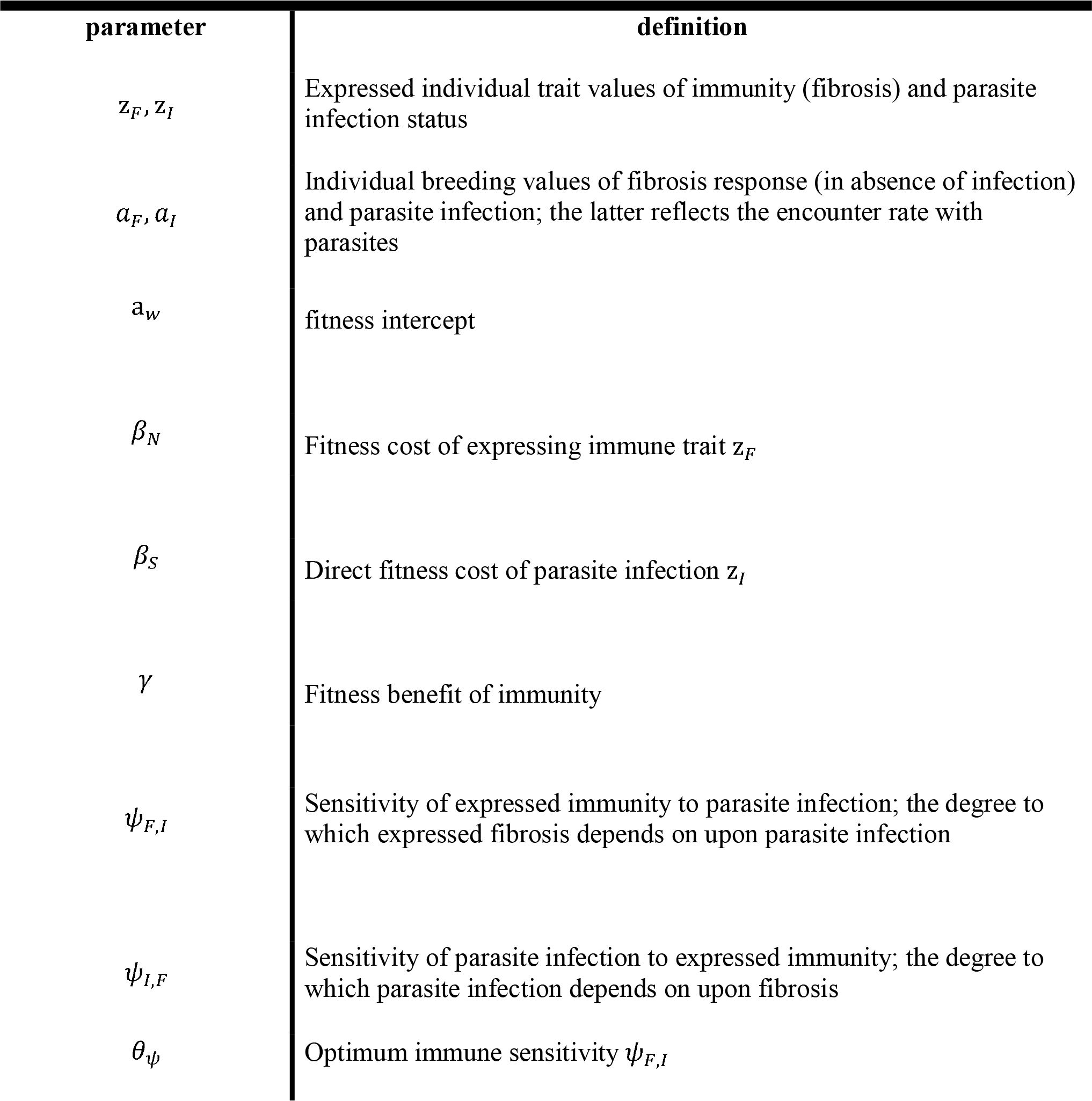
Definition of model parameters

| parameter | definition |
| --- | --- |
| $z_F, z_I$ | Expressed individual trait values of immunity (fibrosis) and parasite infection status |
| $a_F, a_I$ | Individual breeding values of fibrosis response (in absence of infection) and parasite infection; the latter reflects the encounter rate with parasites |
| $a_w$ | fitness intercept |
| $\beta_N$ | Fitness cost of expressing immune trait $z_F$ |
| $\beta_S$ | Direct fitness cost of parasite infection $z_I$ |
| $\gamma$ | Fitness benefit of immunity |
| $\psi_{F,I}$ | Sensitivity of expressed immunity to parasite infection; the degree to which expressed fibrosis depends on upon parasite infection |
| $\psi_{I,F}$ | Sensitivity of parasite infection to expressed immunity; the degree to which parasite infection depends on upon fibrosis |
| $\theta_\psi$ | Optimum immune sensitivity $\psi_{F,I}$ |

Importantly, the two traits in this model *z_F_* and *z_I_* are not expected to be expressed independently, and in fact are measurable outcomes of underling latent trait(s). Fibrosis is induced by parasite exposure, but in turn it acts to reduce the probability of successful infection, *z_I_*. Thus fibrosis, *z_F_*, is a trait whose expression is determined not only by host genotype but also by parasite exposure, an environmental effect. Infection, *z_I_*, is a trait determined both by parasite encounter rates as well as the mitigating effects of fibrosis response. We define trait expression as

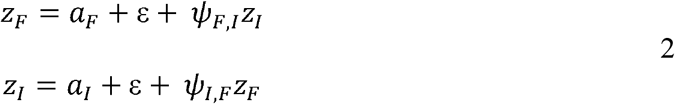

where *α_F_* and *α_I_* are intercepts that describe the additive breeding value for the trait. For fibrosis, this represents fibrotic expression in the absence of exposure to parasites, and may typically be close to zero. For parasite infection, *α_I_* represents exposure rate in the absence of immune response, or the probability of exposure to parasites, and thus is an ecological variable determined by diet and habitat choice.

The terms *ψ* represent the degree to which trait expression is influenced by another trait, in this case infection status or fibrosis. Thus *ψ_F,I_* is a latent trait describing the degree to which fibrosis is induced in response to a parasite exposure, and is thus the key variable of interest in our model, whose evolution we seek to predict. We borrow our notation here from the literature on indirect genetic effects (Moore et al. 1997, McGlothlin et al. 2010), because these effects of immune response on infection repression and infection induction of immune response can be considered a special case of an indirect genetic effect (we develop IGE models of host-parasite coevolution more extensively in another paper). We can now define each trait explicitly in terms of the other,

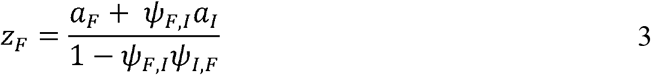

with a similar equation for *z_I_*. We note that in our model of antagonistic interactions between immune response and parasite infection, we expect *ψ_F,I_* to range from 0 to 1 and for *ψ_I,F_* to range from 0 to −1.

We can now analyze this model to determine how selection acts on the latent immune response trait *ψ_F,I_*. Substituting equations 2 and 3 into 1 and differentiating with respect to *ψ_F,I_* yields a selection gradient:

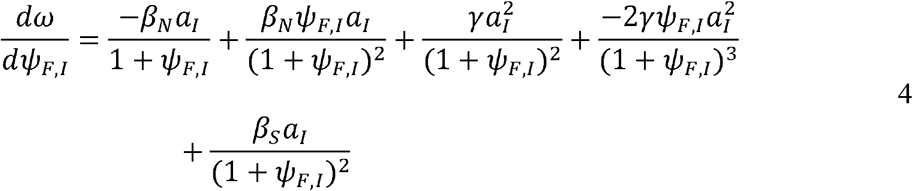

assuming for simplicity that *α_F_* = 0 and *ψ_I,F_* = −1. Equation 4 has three roots representing evolutionary equilibria. One is at *α_I_* = 0, an ecological scenario where parasites are absent or never encountered. Another is at *γ* = 0 and *β_N_* = *β_S_*, where there is no benefit of fibrosis and costs of immune response equal costs of parasite infection. Finally, the relevant root for evolution of *ψ* is

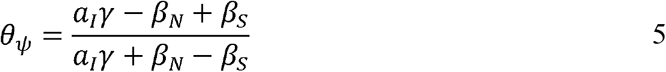

which holds for *α_I_γ* + *β_N_* ≠ *β_S_* and *α_I_γ* ≠ 0, and is a local optimum (see Appendix).

Equation 5 describes the location of the optimum immune response sensitivity as a function of the costs and benefits of immune response and parasite infection. Immediately notable is that increases in parasite prevalence, captured by the intercept *α_I_* lead to an increase in the optimum sensitivity (Figure 7A). However, regardless of the magnitude of *α_I_* when the natural selection cost of fibrosis, *β_N_*, is less than or equal to the cost of infection *β_S_*, the optimum is for high sensitivity (Figure 7B, C, E). Intriguingly, we can also see that immune response can evolve even when the natural selection costs of fibrosis outweigh the costs of carrying an infection (Figure 7D), if the prevalence of parasites is high enough and the costs of infection are in part mitigated by the interaction term *γ*.

**Figure 7.**
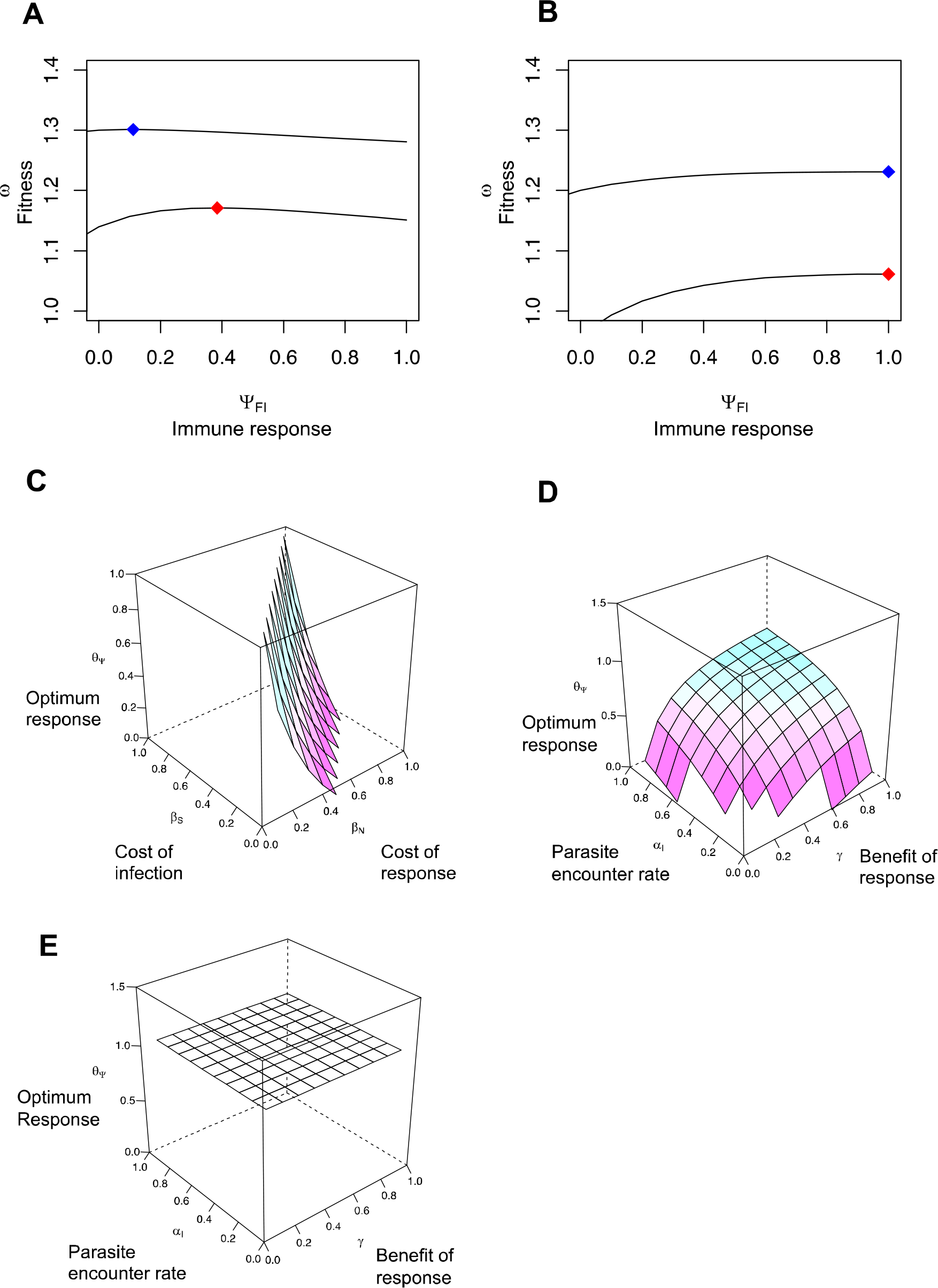
A model of immune response evolution. Panels A and B show fitness as a function of immune response sensitivity *ψ_F,I_* under two parasite encounter rates, *α_I_* = 0.5 (blue optimum, *θ_ψ_*) and *α_I_* = 0.9 (red optimum, *θ_ψ_*), where costs of fibrosis exceed costs of infection (A) and where costs of fibrosis equal costs of infection (B). Panel C illustrates the optimum as a function of costs of fibrosis and costs of infection, holding *γ* and *α_I_* constant; values where *β_N_* < *β_S_* all exceed unity. When *β_N_* > *β_S_*, the optimum is determined by the mitigating effects (*γ*) of fibrosis on fitness costs of parasite infection, as well as parasite encounter rate *α_I_* (Panel D). When *β_N_* ≥ *β_S_*, these mitigating effects and encounter rates play little role in the position of the optimum (Panel E).

The main conclusion from this model is that although optimum immune sensitivity can indeed be governed by encounter rate, when costs and benefits of infection and immune response are high these encounter rates play less of a role in determining the optimum immune response. We find evidence of sex and population differences in parasite encounter rates, *α_I_*, based on differences in worm infection rates across populations and sexes (Figure 1) and sex and population differences in diet (Figure 5; see also Bolnick and Ballare 2020). Yet we find no evidence of sex differences in fibrotic immune response in the wild, and past work in the lab confirms this for Roselle lake and other Vancouver Island populations (Hund et al. 2020). Moreover, QTL mapping of fibrosis response to experimental cestode infection in the lab revealed no sex effect on fibrosis (Weber et al. in prep). At present, we lack data on lifetime reproductive success (in addition to a way to estimate *α_I_* directly), and so cannot confront our model directly with the data at hand. But, this model provides guidance for key parameters that should be measured in the future. And, the data we do have shows that in both populations and sexes there are measurable fitness costs of infection and fibrosis. We can get a better understanding of these costs by re-fitting linear models with component fitness as a predictor and binary infection status, fibrosis status, and their interaction as fixed effects. From these regression coefficient estimates, we can take the linear terms and multiply them by −1 and use the raw interaction term to obtain estimates of the parameters in Equation 1. We assume normality for the sake of parameter estimation and comparison. Corresponding estimates of *γ*, *β_N_*, and *β*_S_ for each subpopulation sampled are provided in Table 2.

**Table 2.**
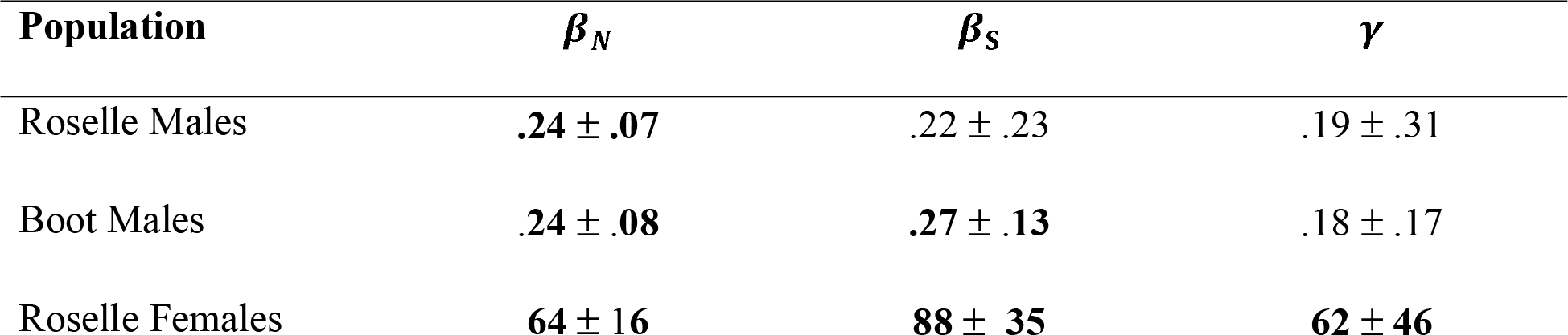
Estimates of parameters in Equation 1. Bold indicates statistical significance, with the caveat that response variables (component fitness) were not normally distributed. ± 1 Standard Error

We can reach two key conclusions from these parameter estimates. First, costs of fibrosis and infection (*β_N_*, and *β*_S_) are always higher than the mitigating effects of *γ*. Second, *β_N_*, and *β*_S_ are either similar in magnitude (males in Boot and Roselle) and/or *β_N_* < *β*_S_ (Roselle females, Roselle males). Thus, these values are consistent with a high optimum for *ψ_F,I_* and relatively low sensitivity of that optimum to parasite encounter rates, *α_I_*. Thus, our model and data suggest the sexes can be expected to exhibit similar optimum immune responses when the costs and benefits of immune response are high and sexually concordant, even if exposure rates to the parasites are sexually dimorphic due to sex differences in diet or habitat use.

## Discussion

Numerous evolutionary models suggest that the strength or sensitivity of an immune response should be subject to net-stabilizing selection, rather than persistent directional selection, to balance the costs and benefits of immunity (Sheldon and Verhulst 1996, Lochmiller and Deerenberg 2000, Zuk and Stoehr 2002, Graham 2013). The costs of immunity are well documented and include increased energetic expenditure and decreased resource acquisition (Houston et al. 2007), increased vulnerability to predators (Navarro et al. 2004), loss of social status (Stockmaier et al. 2020), and the increased risk of immune pathology (Goldzmid and Trinchieri 2012). These costs combine with the benefits of immune response to costly infections to determine the optimum immune response. Thus, quantification of the costs and benefits of immune response is key to understanding the evolutionary origins of variation in parasite infection rates and host immune evolution observed across populations and sexes in the wild.

Here, we have presented a rare instance of an immune response with measurable benefits and costs in the wild. In two lakes on Vancouver Island, we found that peritoneal fibrosis is associated with reduced cestode infection, both in terms of infection probability and parasite growth when infected. However, we find that this immune response comes at substantial component fitness costs for both sexes. Fibrosis is associated with a reduction in prey consumption. In both lakes, fibrotic males are less likely to be reproductively successful, indicating consistent selection against this immune response trait. Similar reproductive costs are observed in females, with substantial reduction in ovary mass in individuals expressing fibrosis (in the one lake with enough infected females to test this trend). We show that these shared fitness costs act to align multivariate selection across males and females, which would otherwise entail sexually-antagonistic selection on trophic morphology.

Consistent with our results, recent laboratory experimental infection studies demonstrate that exposure to cestode antigens induces fibrosis (Hund et al. 2020) which in turn suppresses cestode growth and viability (Weber et al. in prep). In these laboratory assays, the sexes were equally prone to fibrosis following an immune challenge. Although males and females initiate similar levels of fibrosis following a controlled immune challenge, their risk of cestode infection should be unequal. In many lake populations of stickleback females feed on more benthic prey than males do, though in some lakes this diet dimorphism is absent or reversed. *S.solidus* is acquired by eating limnetic prey (cyclopoid copepods), so diet dimorphism should, and does, generate sex differences in cestode encounter and infection rates (Reimchen and Nosil 2001, Stutz et al. 2014). At face value, this presents a conundrum because substantial costs of fibrosis, coupled with sex differences in parasite encounter rates, might be intuited to result in evolution of sex differences in immune sensitivity to cestode infection. Our analysis of a simple model of immune response evolution, however, indicates that optimum immune response can be surprisingly insensitive to parasite encounter rates when costs of both immune response and parasite infection are high. This finding can reconcile our results, and also indicates that interpreting infection rates alone as a measure of immune activity (as is often done in the case of sex differences in parasite loads) may be problematic (Poulin 1996). Our model also provides a glimpse into the complex forms of selection that may inherently act on indirect genetic effects, which are typically (Moore et al. 1997, McGlothlin et al. 2010) assumed to be fixed population parameters in theoretical models.

Cestode prevalence varies widely across lakes in Western Canada (Stutz et al. 2014, Weber et al. 2017b), and some stickleback populations do not exhibit fibrotic response to cestode infection (Weber et al. in prep, Hund et al. 2020). Our results indicate that variation in the costs and benefits of fibrosis across populations could explain this variation in stickleback immune response. When the fitness cost of fibrosis exceeds the costs associated with cestode infection, the parameter space where fibrosis response to parasite exposure is expected to evolve in our model is limited. Our data from two populations indicate that the magnitude of these fitness costs are very similar, and even slightly higher for fibrosis in males from Roselle lake. Thus, even slight variation in the relative costs of fibrosis across populations could result in selection against fibrosis response in some lakes. Variation in the cost of fibrosis could arise from variation in the strength of competition for mating opportunities in males and/or variation in the strength of selection on female reproductive output, both of which (Boughman 2007, Baker et al. 2008, Heins and Baker 2014) are likely to vary across stickleback populations.

Sexual conflict theory indicates that sexually concordant fitness effects can mask the signature of sexual antagonism. This occurs if concordant effects are strong enough to override otherwise-conflicting selection acting upon the sexes, thus acting to align net selection across males and females. One way these effects are often envisaged is as a displacement from sex-specific optima, where maladaptation pushes both sexes away from their respective fitness peaks, aligning selection on the sexes transiently until the population adapts to its new environment and conflict again ensues (Long et al. 2012, Connallon 2015, Connallon and Hall 2016). Our analysis demonstrates a slightly different manifestation of how concordant fitness effects can change the nature of fitness variance across the sexes. In our study, we show that concordant fitness effects of a shared trait – fibrosis – act to align otherwise antagonistic selection acting through morphology when both suites of traits are included in a multivariate analysis. Selection is antagonistic and concordant in different dimensions of multivariate trait space, with the direction of net multivariate selection largely aligned due to the strong concordant fitness effects of fibrosis that swamp out antagonistic effects of morphological shape. Thus, our work illustrates how specific traits can contribute to or mask sexually antagonistic fitness variance. To the extent that immune response optima are shared across the sexes, such effects could lead to the appearance of pervasive sexual conflict when they are not measured, creating a ‘hidden-trait’ problem (Morrissey et al. 2010) that could contribute to the appearance (Cox and Calsbeek 2009) of widespread unresolved sexual conflict in the wild.

By surveying component-fitness costs and benefits of parasite infection and immune response, we show that relative costs of infection and immune response can be similar across males and females despite acting through different fitness components across the sexes. Our simple analytical model of immune response evolution indicates that similar male and female immune optima may be a common feature across a range of ecological scenarios, especially when fitness costs of immune response and parasite infection are high. Thus, our work illustrates how shared fitness costs in host-pathogen interactions can lead to shared evolutionary optima for male and female host immunity.

## Acknowledgements

We are grateful to Will Shim and Jacqueline Salguero for their help in sampling fish, as well as Nathalie Steinel and Jesse Weber for logistical support in the field. Lauren Fuess and Amanda Hund provided help on scoring fibrosis. We thank Katherine Lewkowicz and Megan Braat for assistance with the dissections. A. Hund, Milan Vrtilek, and Foen Peng provided valuable feedback on the manuscript. Funding was provided by the University of Connecticut to DIB, and NIAID Grant 1R01AI123659-01A1 to DIB.

## Appendix

We can confirm that the root of equation 4 given in equation 5 is a local optimum by solving for the second derivative at the root,

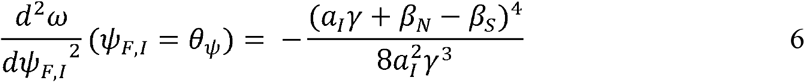

Which will be negative for all *γ* > 0.

**Table S1.** Fish numbers sampled

| Boot Lake |  |  |
| --- | --- | --- |
|  | Infected | Uninfected |
| Nesting Male | 9 | 43 |
| Trapped Male | 32 | 70 |
| Trapped Female | 24 | 6 |
| Roselle Lake |  |  |
|  | Infected | Uninfected |
| Nesting Male | 2 | 48 |
| Trapped Male | 7 | 99 |
| Trapped Female | 7 | 64 |

